# Role of socioeconomic factors and interkingdom crosstalk in the dental plaque microbiome in early childhood caries

**DOI:** 10.1101/2024.03.12.584708

**Authors:** Mohd Wasif Khan, Vivianne Cruz de Jesus, Betty-Anne Mittermuller, Shaan Sareen, Victor Lee, Robert J. Schroth, Pingzhao Hu, Prashen Chelikani

## Abstract

Early childhood caries (ECC) is influenced by microbial and host factors, including social, behavioral, and oral health. In this cross-sectional study, we analyzed interkingdom dynamics in the dental plaque microbiome and its association with host variables. The samples collected from the preschool children underwent 16S rRNA and ITS1 rRNA gene sequencing. The questionnaire data were analyzed for social determinants of oral health. The results indicated a significant enrichment of *Streptococcus mutans* and *Candida dubliniensis* in ECC samples, in contrast to *Neisseria oralis* in caries-free children. Our interkingdom correlation analysis revealed that *Candida dubliniensis* was strongly correlated with both *Neisseria bacilliformis* and *Prevotella veroralis* in ECC. Additionally, ECC showed significant associations with host variables, including oral health status, age, place of residence, and mode of childbirth. This study provides empirical evidence associating the oral microbiome with socioeconomic and behavioral factors in relation to ECC, offering insights for developing targeted prevention strategies.

**HIGHLIGHTS:**

- Characterized interkingdom association between cariogenic species of genus Neisseria and Candida
- Both bacterial and fungal species are important for caries status prediction using artificial intelligence
- Socioeconomic index is associated with caries status and caries-associated microbial markers

## INTRODUCTION

Early childhood caries (ECC) is a condition involving tooth decay in the primary dentition in children under 72 months of age. The prevalence of ECC is a major public health concern worldwide, affecting nearly half of all children globally (El Tantawi et al., 2018; Uribe et al., 2021) In Canada, the prevalence of ECC in some disadvantaged communities can be as high as 98.5% (Pierce et al., 2019). This high prevalence imposes significant socioeconomic and psychological burdens on families and society (Bencze et al., 2021; Carvalho et al., 2018). The link between ECC and future caries experiences in permanent dentition highlights its importance as a critical factor in long-term oral health (Jordan et al., 2016; Tsai et al., 2023). Furthermore, ECC can increase the risk of other dental diseases, malocclusion, negative oral health related quality of life (OHRQoL), and nutritional status (Davidson et al., 2016; Khan et al., 2023; J. Lee et al., 2022; Schroth et al., 2022).

The oral cavity is known to harbor more than 700 species of bacteria and 100 species of fungi (Diaz & Dongari-Bagtzoglou, 2021; Escapa et al., 2018; Peters et al., 2017). Acidogenic and aciduric microorganisms are crucial in the pathogenesis of dental caries and form an integral part of dental plaque biofilms identified in caries (Alam et al., 2000; X. Chen et al., 2020; Struzycka, 2014). Dysbiosis in microbial composition, characterized by an overabundance of acidogenic microbes, increases the risk of caries (J. Chen et al., 2021; Grier et al., 2020).

Constant exposure to acidic conditions leads to the demineralization of tooth enamel (Marsh & Nyvad, 2008; Takahashi & Nyvad, 2011). This effect can be partially reversed by using fluoride, which alters the dental plaque microenvironment and promotes remineralization of the enamel (Zhang et al., 2022). Host variables, encompassing various demographic, socioeconomic, and behavioral factors, also influence the development of caries (Arora et al., 2011; Imes et al., 2021; Pierce et al., 2019). Therefore, researchers are increasingly interested in investigating the comprehensive role of these risk factors in ECC, such as diet, fluoride exposure, limited access to care, and poor oral hygiene (Adler et al., 2021; Ganesh et al., 2020; Handsley-Davis et al., 2021; Kahharova et al., 2023; Schroth et al., 2009).

Bacteria and fungi, present on all human sites in contact with the external environment, exhibit a diverse range of interactions. These interactions, both symbiotic and competitive, play a crucial role in maintaining the microenvironment (Barbosa et al., 2016; He et al., 2017). A similar trend is observed in ECC-associated bacteria and fungi, where *Streptococcus mutans* and opportunistic *Candida albicans* can exhibit both synergistic and antagonistic behaviors in ECC (Lu et al., 2023). These interactions can affect the host through pathogenesis induced by ecological shifts (Balakrishnan et al., 2021). However, previous studies have generally limited their focus to interkingdom interactions at the genus level or specific microbial species (Kim et al., 2021; Krzyściak et al., 2017; Tu et al., 2022). Therefore, further investigations are necessary to enhance our understanding of interkingdom microbial signaling and their interactions with host risk factors in caries development.

Using a cross-sectional design, we analyzed dental plaque samples from preschool children (n = 538) by advanced sequencing techniques, statistical, and machine learning analyses. The objectives of this study were twofold: first, to enhance our understanding of the role of social and behavioral variables associated with ECC, and second, to identify the interkingdom interactions between bacteria and fungi within the dental plaque microbiome associated with ECC. Our study provides a comprehensive analysis of the ECC microbiome, considering a broad spectrum of host variables, including socioeconomic and behavioral factors. We expect that our findings based on these variables will enhance the screening of young children at a high risk of caries and the development of informed strategies for intervention.

## METHODOLOGY

### Participant recruitment and sample collection

This study enrolled 553 participants, who were recruited between December 2017 and July 2022 at the Misericordia Health Centre (MHC), Children’s Hospital Research Institute of Manitoba (CHRIM), and various community dental clinics in Winnipeg, Manitoba, Canada. Among the participants, 538 children were under the age of 72 months. Children aged > 72 months or missing age information (n = 15) were excluded from the study. The remaining participants included 226 caries-free (CF) children and 312 children with ECC, implementing a cross-sectional study design. The dental examination was performed by experienced dentists to determine the CF and ECC status. From these participants, supragingival plaque samples were collected from all tooth surfaces (Agnello et al., 2017). Children on any antibiotic and who did not meet the case definition for ECC were not selected for this study (American Academy of Pediatric Dentistry, 2003). At MHC, the samples were collected before the scheduled surgery, and for other locations, the samples were obtained during standard oral examinations by a dentist or research staff. The participants’ information was collected as metadata through a questionnaire completed by parents or caregivers who also provided written informed consent. Participant recruitment, sample collection, and information collection for this study were approved by the University of Manitoba’s Health Research Ethics Board (HS22388 – H2018:472). This study adhered to the checklist provided by the STROBE guidelines for conducting cross-sectional studies (**Supplementary Table S1**). The methodology for dental plaque sample collection and DNA extraction was consistent with the procedures described in our previous studies (de Jesus et al., 2021, 2022). To estimate the statistical power of our microbiome study, an online tool “micropower” was used for supragingival plaque at a significance level of 0.05, while maintaining other parameters at their default settings (Kelly et al., 2015).

### Amplicon sequencing

Library preparation and sequencing were performed using either MiSeq or NovaSeq6000 instruments (Illumina Inc., San Diego, CA, USA), employing paired-end (250 × 2 bp in length) sequencing technology at the Genome Quebec Innovation Center (Montreal, Canada), using amplicon sequencing targeting the V4 hypervariable region of bacterial 16S rRNA and fungal ITS1 (internal transcribed spacer 1) spacer DNA. Primers for the 16S rRNA V4 region were 515F (5′-GTGCCAGCMGCCGCGGTAA-3′) and 806R (5′-GGACTACHVGGGTWTCTAAT-3′).

Primers targeting the ITS1 region were ITS1-30 (5′-GTCCCTGCCCTTTGTACACA-3′) and ITS1-217 (5′-TTTCGCTGCGTTCTTCATCG-3′) (de Jesus et al., 2020; Usyk et al., 2017).

### Sequencing data processing and taxonomic assignment

The sequencing reads received from the sequencing center were provided as demultiplexed, barcode-removed, and paired-end FASTQ files. These reads were processed using QIIME2 (version 2022.11) to create separate amplicon sequence variant (ASV) tables for 16S and ITS1 reads (Bolyen et al., 2019). For the 16S sequencing data, quality trimming, filtering, removal of chimeric reads, and merging were performed using DADA2 within the QIIME2. The read trimming length for each sequencing batch was optimized using different combinations of forward and reverse reads, and the best combination was selected for final trimming. For ITS data, the Q2-ITSxpress QIIME2 plugin was used to trim the conserved regions around ITS1 before applying DADA2 for merging and chimera removal (Bengtsson-Palme et al., 2013; Rivers et al., 2018). The Human Oral Microbiome Database (HOMD, version 15.23) and UNITE database, which uses dynamic clustering thresholds (version 9; QIIME release), were employed for taxonomic assignment of bacterial and fungal ASVs, respectively. Separate ASV tables consisting of the abundance at each taxonomic level were generated for bacteria and fungi. The ASV tables were filtered to retain features that exhibited a minimum prevalence of 5% across all the samples.

### Microbiome community and network analysis

All community analyses were performed using the phyloseq object in R package “phyloseq” (version 1.40.0). For alpha diversity, the Chao1, Shannon, and Simpson diversity indices were used. Beta diversity was calculated using the adonis2 function in the “vegan” package (version 2.6-4). For the plotting of the most abundant species “microeco” (version 1.4.0) options were considered. All figures were generated in the “ggplot2” (version 3.4.3) library in R.

To identify the differentially abundant species between CF and ECC, association analysis with a linear model was implemented using MaAsLin2 (version 1.14.1) in R (Mallick et al., 2021).

MaAsLin2 is a statistical model designed to identify the associations between microbial taxa and clinical metadata. It employs general linear models and incorporates methods for data normalization and transformations to identify multivariable associations, while controlling for the false discovery rate. In our application of MaAsLin2, the “LM” method was utilized on center log-ratio (CLR) transformed abundance data as the response variable and ECC status as the independent variable. In this analysis age, sex, and place of residence were considered as confounding variables along with the Benjamini-Hochberg (BH) method to control for false discovery rates. To further minimize the number of false positives, associations based on a q-value (adjusted p-value) threshold of less than 0.01 were considered significant.

For the correlation network analysis, the species identified by MaAsLin2 were used to generate a network plot illustrating the associations between species using the “NetCoMi” package (version 1.1.0) in R. First, network associations for the CF and ECC groups were obtained separately, employing the association measure “SPRING”. Subsequently, a differential network was generated from these two groups using “discordant” as the differential network method.

### Classification of CF and ECC samples

To discriminate between CF and ECC samples, machine learning (ML) classifiers were constructed in R using the “tidymodels” package (version 1.1.1). To perform the classification between CF and ECC samples, we applied commonly used ML methods (Topçuoğlu et al., 2021; Wirbel et al., 2020). We tested five ML classifiers: LASSO logistic regression, ridge logistic regression, Support Vector Machine (SVM), Random Forest (RF), and LightGBM. A stratified 5-fold cross-validation approach with 10 repeats was employed. The features used in these classifiers were the CLR normalized abundance data for 16S and ITS sequences.

For the lasso and ridge logistic regression, the ’glmnet’ package was utilized for L1 and L2 regularization, respectively, with fine-tuning of the loss-function hyperparameter. In the SVM model, the cost and degree hyperparameters were fine-tuned, with the other parameters set to their default values. For the RF model, the number of trees, the minimum node size, and the number of variables at each split were tuned. In the LightGBM model, the tree depth and learning rate were fine-tuned along with the hyperparameters used in the RF model. To ensure a comprehensive analysis, a grid size of 50 was used for tuning the hyperparameters in all selected classifiers. The performance of each method was compared using the Area Under the Receiver Operating Characteristic Curve (AUROC) and Precision-Recall Curve (AUPRC) metrics.

### Analysis of socioeconomic and behavioral variables

Demographic, lifestyle, socioeconomic, and behavioral variables were collected using a comprehensive questionnaire obtained from parents or caregivers at the time of participant recruitment. These variables will be referred to as ’host variables’ and include age, sex, feeding, and oral hygiene habits, among others (**Table 1)**. Residential information using the postal codes of the participants was utilized for rural-urban designation as well as to determine the Socio- Economic Factor Index (SEFI) score, social deprivation score, and material deprivation score (Fransoo et al., 2013). These data were obtained from the Manitoba Centre for Health Policy (MCHP) (Metge et al., 2009). The Early Childhood Oral Health Impact Scale (ECOHIS) and NutriSTEP scores were determined based on responses to a standard set of questions (Pahel et al., 2007; Randall Simpson et al., 2008).

**Table 1:** Host variables analyzed in the study.

|  | Variable Name | Type | Description | Encoding |
| --- | --- | --- | --- | --- |
| 1 | ECC Status | Binary | Indicates the dental caries status of the child. | Caries free (0), ECC (1) |
| 2 | Sex | Binary | The biological sex of the child. | Female (0), Male (1) |
| 3 | Vaginal Birth | Binary | Mode of childbirth. | Cesarian (0), Vaginal (1) |
| 4 | Breastfeed Child | Binary | Whether the child was breastfed. | No (0), Yes (1) |
| 5 | Bottlefeed Child | Binary | Whether the child was bottle-fed. | No (0), Yes (1) |
| 6 | Bedtime Snack | Binary | Consumption of snacks by the child at bedtime. | No (0), Yes (1) |
| 7 | Urban_Rural | Binary | The residential setting of the child's home. | Rural (0), Urban (1) |
| 8 | Times Day Brushed | Ordinal | Number of times the child's teeth are brushed per day. | Never (1), Sometimes (2), 1x (3), 2x (4), >2x (5) |
| 9 | Child Dental Health | Ordinal | Parental perception of the child's dental health. | Very good (1), Good (2), Fair (3), Poor (4), Very poor (5) |
| 10 | Age | Continuous | The age of the child. | Measured in months |
| 11 | ECOHIS Total Score | Continuous | Score from the Early Childhood Oral Health Impact Scale measures the oral health-related quality of life (OHRQoL) of preschool children and their families. | Higher ECOHIS score signifies worsening oral health related quality of life. |
| 12 | NutriSTEP Total Score | Continuous | 17 questions about a child's typical food choices, eating behaviors, as well as physical activity, and growth to assess the nutrition risk for preschoolers | A higher score indicates a greater nutritional risk. |
| 13 | SEFI score | Continuous | Derived from Census data that reflects non-medical social determinants of health and is used as a proxy measure of socioeconomic status | Lower scores indicate more favorable conditions |
| 14 | Material deprivation score | Continuous | Calculated from Canadian Census data which reflects the deprivation of wealth, goods, and conveniences | Lower scores indicate less deprivation (better status) |
| 15 | Social deprivation score | Continuous | Calculated from Canadian Census data which reflect the deprivation of relationships among individuals in the family, the workplace, and the community | Lower scores indicate less deprivation (better status) |
| 16 | Fluoride toothpaste | Binary | Use of fluoride toothpaste | No (0), Yes (1) |

Footnote: Binary variables are encoded as 0 or 1 based on the alphabetical order of their categories.

To assess the number of missing entries prior to imputation, the R packages “mice” (version 3.16.0) was used. Missing information in the data was imputed using the default methods in the Multiple Imputation by Chained Equations (MICE) approach for binary, categorical, and numerical data. The data was then checked for correlations among the variables using Spearman’s correlation analysis. Variables were tested for associations with ECC status using both multivariable and univariate approaches. In the multivariable analysis, age, sex, and place of residence were considered as confounding variables.

### Microbiome and socioeconomic variables

The data was analyzed to determine the association between host variables and various microbiome phenotypes. For the alpha diversity index, a non-parametric Spearman correlation test was used. For beta diversity dissimilarities, PERMANOVA was applied using adonis2 on the Bray-Curtis distance with 999 permutations. The p-value for each variable was adjusted using the BH method. All statistical analyses were performed using R version 4.3.2 (R Core Team, 2023). To identify the associations between individual taxa and host variables, MaAsLin2 was used, as described above.

## RESULTS

### Microbiome community analyses

A total of 230 bacterial and 95 fungal species were consisted in the filtered ASV tables obtained from QIIME2 analysis. Based on the number of bacterial species, our sample size would achieve a power of 0.93 at significant level 0.05. The most abundant bacterial species identified in bacterial data were *Haemophilus parainfluenzae, Corynebacterium matruchotii, and Lautropia mirabilis* (**Figure 1A**). The alpha diversity metric, Chao1, revealed significant differences between CF and ECC status suggesting a higher number of total species in ECC samples, however, these differences were not observed in the Shannon and Simpson diversity indices.

**Figure 1:**
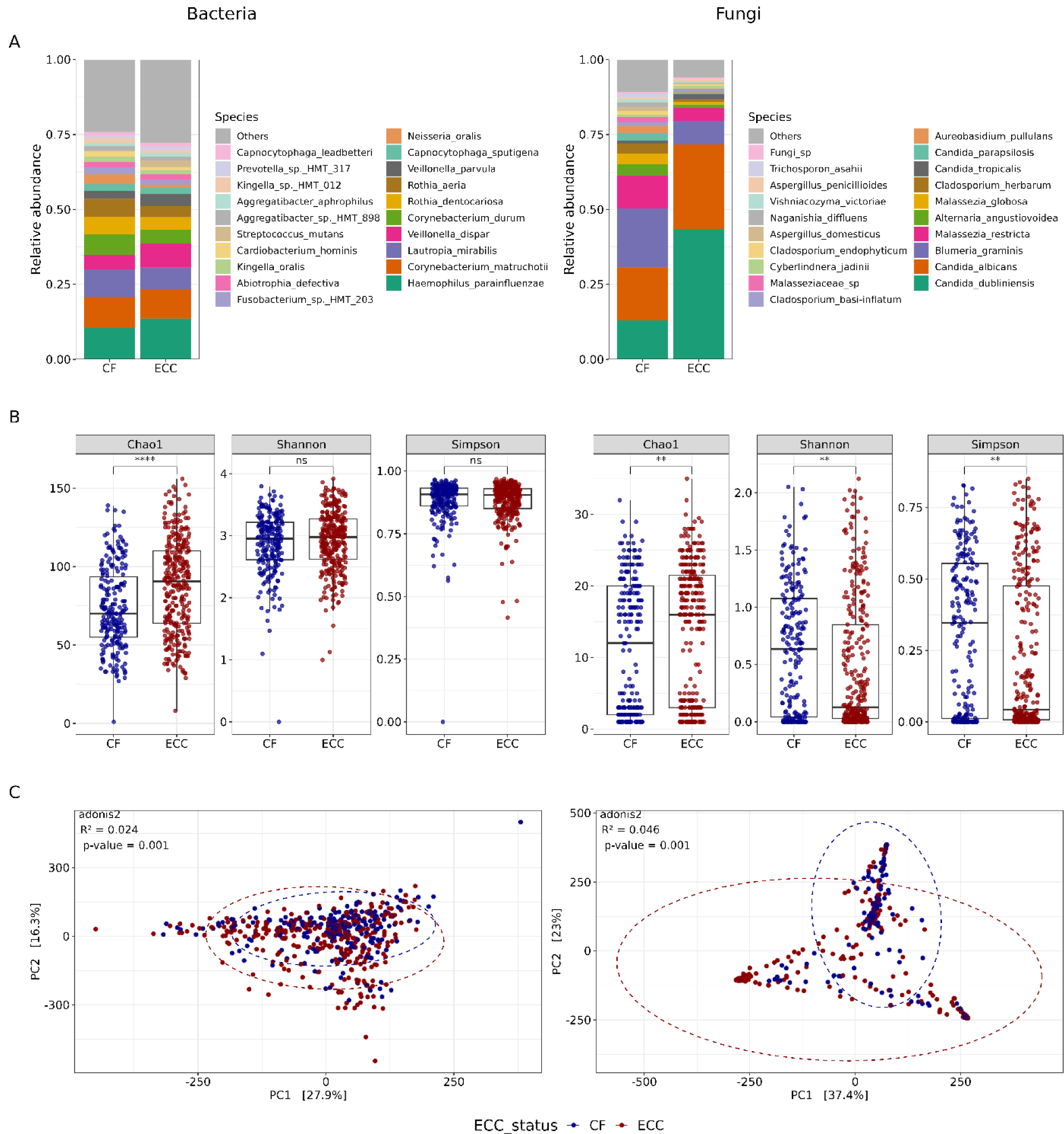
Microbiome composition and alpha and beta diversity in caries-free and early childhood caries. (Left panel for bacteria and right panel represents fungi) (A) Relative abundance of most abundant species. (B) Boxplot for alpha diversity comparison for Chao1, Shannon and Simpson. Each dot represents one sample and boxplot representing median and 1.5 interquartile range. (C) Principal component analysis for beta diversity comparison using adonis2 function.

Chao1 diversity estimates the total number of species in a community based on the number of rare species, that is, species that are present in only a few individuals (**Figure 1B**). Upon comparing the beta diversity differences among the microbial communities, a significant difference was found between CF and ECC (**Figure 1C**).

For fungi, the most common species were *Candida dubliniensis*, *C. albicans,* and *Blumeria graminis* with the former two exhibiting clear differences in their abundance between CF and ECC (**Figure 1A**). The fungal data was found to be significantly different for Chao1, Shannon, and diversity metrics, although the diversity was lower than the corresponding bacterial diversities (**Figure 1B**). The beta diversity in the fungal data was largely similar to the bacterial diversity, showing significant differences between CF and ECC (**Figure 1C**).

### Interkingdom network analysis

To study the correlation among species abundance at both intra and interkingdom levels, we conducted network analyses for ECC and CF samples separately. To visualize ECC-relevant species, we selected only those species found to be associated with ECC status using the MaAsLin2 model. When comparing the networks of the CF samples with those of the ECC samples, more positive connections were observed in the ECC samples than in the CF samples. *Prevotella* species exhibited numerous novel and strong positive associations in ECC samples compared to CF samples. In the differential network plot, *C. dubliniensis* demonstrated positive associations with *Neisseria bacilliformis* and negative associations with *Streptococcus salivarius* and *Prevotella veroralis* (**Figure 2**). Another interkingdom negative correlation in ECC was observed between ECC-associated *C. albicans* and CF-associated *Corynebacterium durum*, *Cladosporium herbarum*, and *Lautrpia mirabilis.* Species exhibiting positive correlations in ECC group included *Selenomonas sputigena*, *P. veroralis*, *Selenomonas flueggei*, *Prevotella salivae*, and *Streptococcus salivarius*. Additional connections involved *S. sputigena*, *P. veroralis*, and *N. bacilliformis*. A third path connecting ECC associated species and originating from *S. sputigena* encompassed *Leptotrichia shahii*, and *Prevotella Olurum*. Another distinct connection, not related to these species, was between *S*. *mutans* and *Scardovia wiggsiae*.

**Figure 2:**
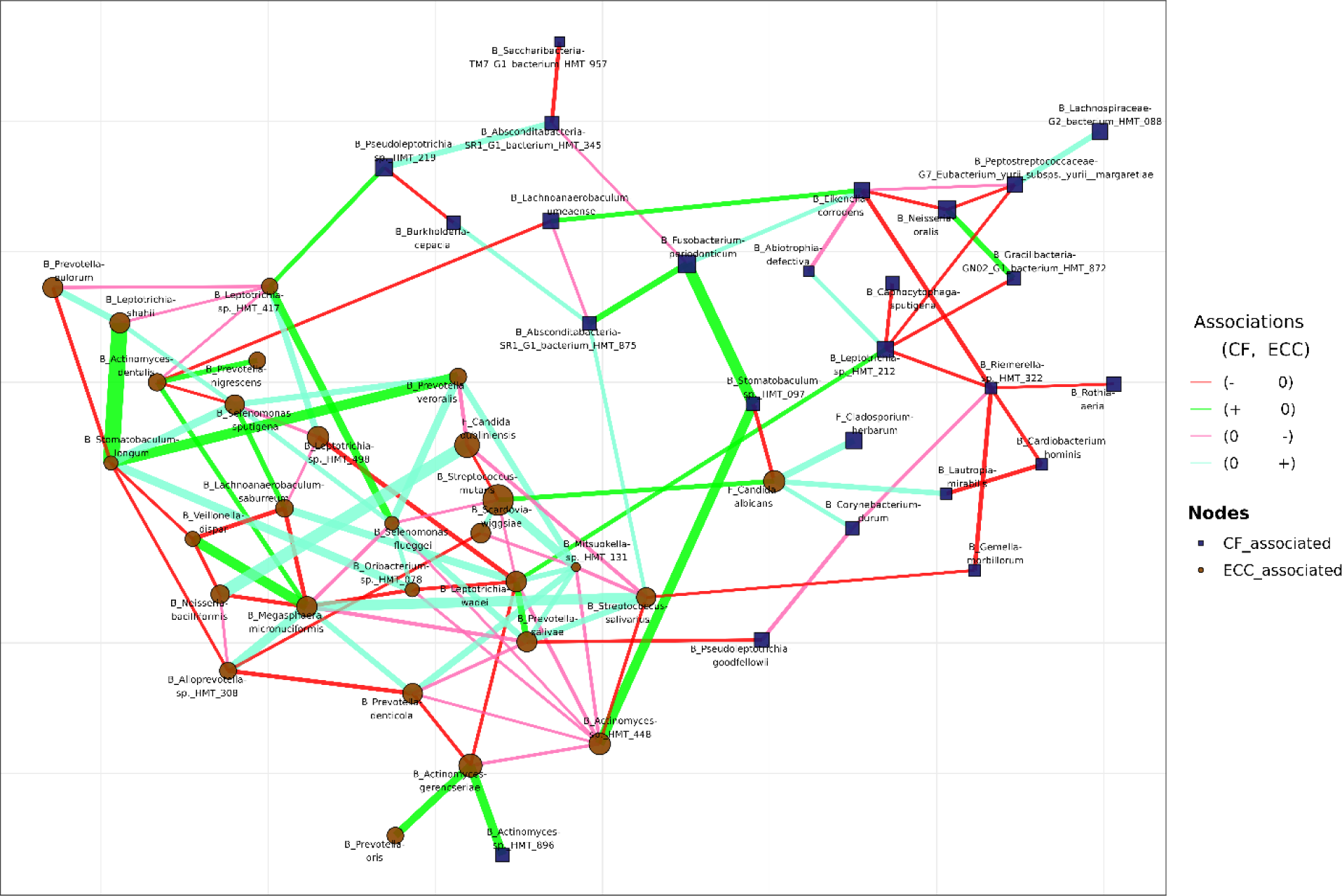
Differential network analysis for CF and ECC samples on combined bacterial and fungal datasets The network plot visualizes the associations between species identified as differentially abundant, highlighting their relevance to CF and ECC conditions. Each node represents one species, and the size of the node is proportional to its coefficient value, indicating the strength of association with either CF or ECC conditions. Edges between nodes signify the interactions from the differential matrix, with each edge type categorized by the nature of interaction within CF or ECC samples: ’0’ denotes no correlation, ’-’ indicates a negative correlation, and ’+’ signifies a positive correlation between two species. To limit the edge connections, only the interactions involving species identified as differentially abundant through MaAsLin2 analysis are included.

Interestingly, *C. albicans* was positively associated with S*. mutans* in CF samples, this association was absent in ECC samples while associations between *C. dubliniensis* and *S*. *mutans* were negative in CF which shifted to no interactions in ECC. The differential network revealed that numerous associations between the CF and ECC samples changed from a state of no interaction to either positive or negative. However, it is noteworthy that we did not identify any instances of drastic shifts in which associations changed from a negative to positive state, or vice versa. Moreover, ECC associated species appeared more dynamic, exhibiting a higher number of both positive and negative associations than those associated with CF.

### Classification of caries status using microbiome data

We evaluated the performance of the classification model separately for bacterial and fungal data, and then on the combined dataset. Among the five machine learning (ML) classifiers selected for the classification of CF and ECC, the RF algorithm emerged as the most effective for bacteria and the combined data of bacteria and fungi, achieving AUROC and AUPRC values of 0.92 and 0.93, respectively (**Figure 3A**). The performance of RF and LightGBM was comparable for fungi, with AUROC and AUPRC values of 0.85 and 0.89, respectively. Other classifiers, such as Lasso, Ridge, and SVM, displayed a performance similar to that of RF for bacteria; however, they did not perform as well as RF or LightGBM for fungi. Except for LightGBM, the overall performance of these classifiers on bacterial data was slightly better than that on fungal data, while the performance on combined data did not improve the results further. The AUPRC values remained consistent for both bacteria and combined data; however, they were notably higher for fungi than their respective AUROC values. Further investigation into the RF model, focusing on combined data for feature importance based on the “permutations” measure, revealed that the species identified as most significant were *S. mutans* and *C. dubliniensis*, with *S. mutans* being significantly more important than any other species (**Figure 3B**). The other top species that appeared in the RF model were *N. bacilliformis*, *Prevotella denticola*, *C. albicans*, and *S. wiggsiae*.

**Figure 3:**
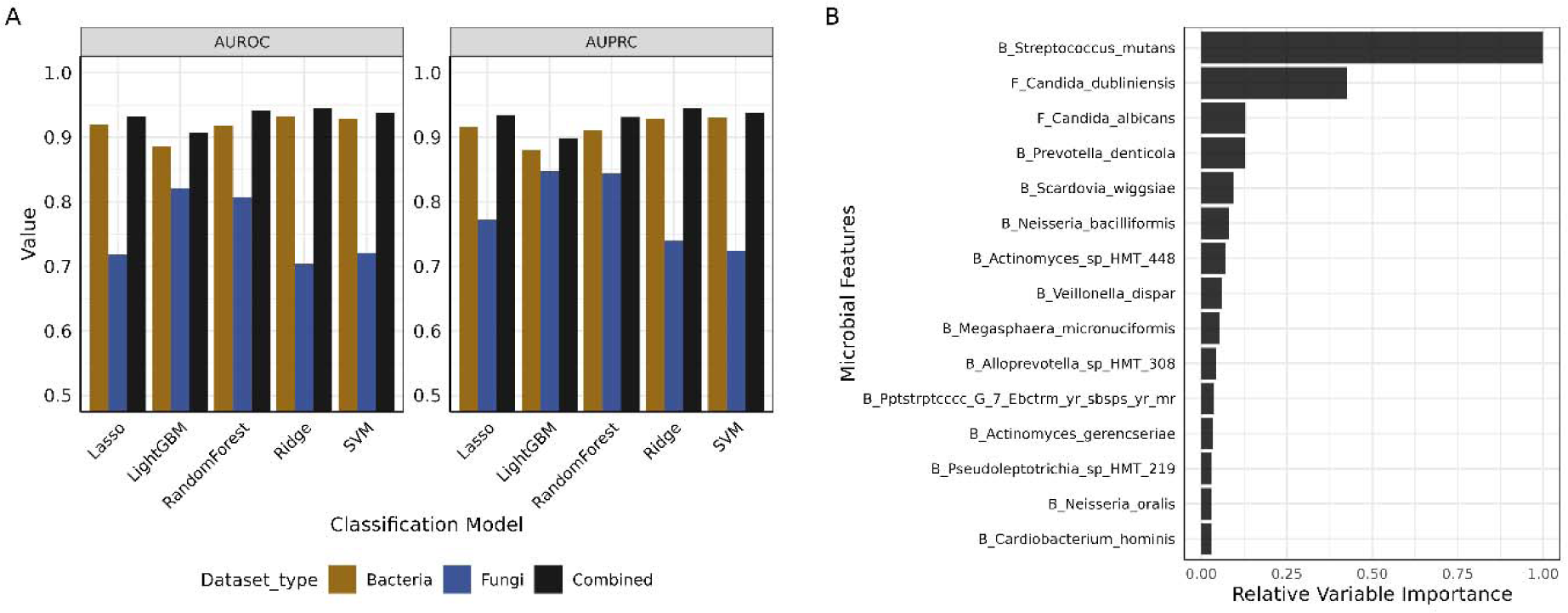
Machine learning performance for the classification of ECC and CF status (A) Barplot for Area Under the Receiver Operating Characteristic Curve (AUROC) and the Area Under the Precision-Recall Curve (AUPRC) values of five machine learning classifiers for the classification of ECC and CF samples, evaluated for the combined dataset and for bacterial and fungal datasets separately. (B) Barplot for the variable importance derived from Random Forest model on combined bacterial and fungal datasets.

### Association of host variables with caries status

A total of 16 host variables from 538 participants were included in the analysis. A summary of these variables is presented in **Table 2**. The ECC status, age, and sex information were not missing from any of the samples, on the other hand, NutriSTEP and ECOHIS scores were not available in 79 and 80 entries, respectively. The distributions of these variables after using MICE imputation for the missing values are shown in **Figure S1.** Spearman correlation analysis revealed a 0.95 coefficient value between the material deprivation score and the SEFI score (**Figure 4A**); therefore, only the SEFI score was used in all subsequent analyses. ECC status, ECOHIS score, and child dental health were also correlated with a correlation coefficient of > 0.65. A negative correlation was observed between the ECC status and urban residence. Our multivariable logistic regression analysis identified a strong association between ECC and other host variables such as child dental health, bedtime snacking, and residency status (**Figure 4B**). Of these variables, child dental health, fluoride toothpaste, vaginal birth, bedtime snacking, age, and ECOHIS score were positively associated with ECC, whereas social deprivation score exhibited a negative trend, though not statistically significant.

**Figure 4:**
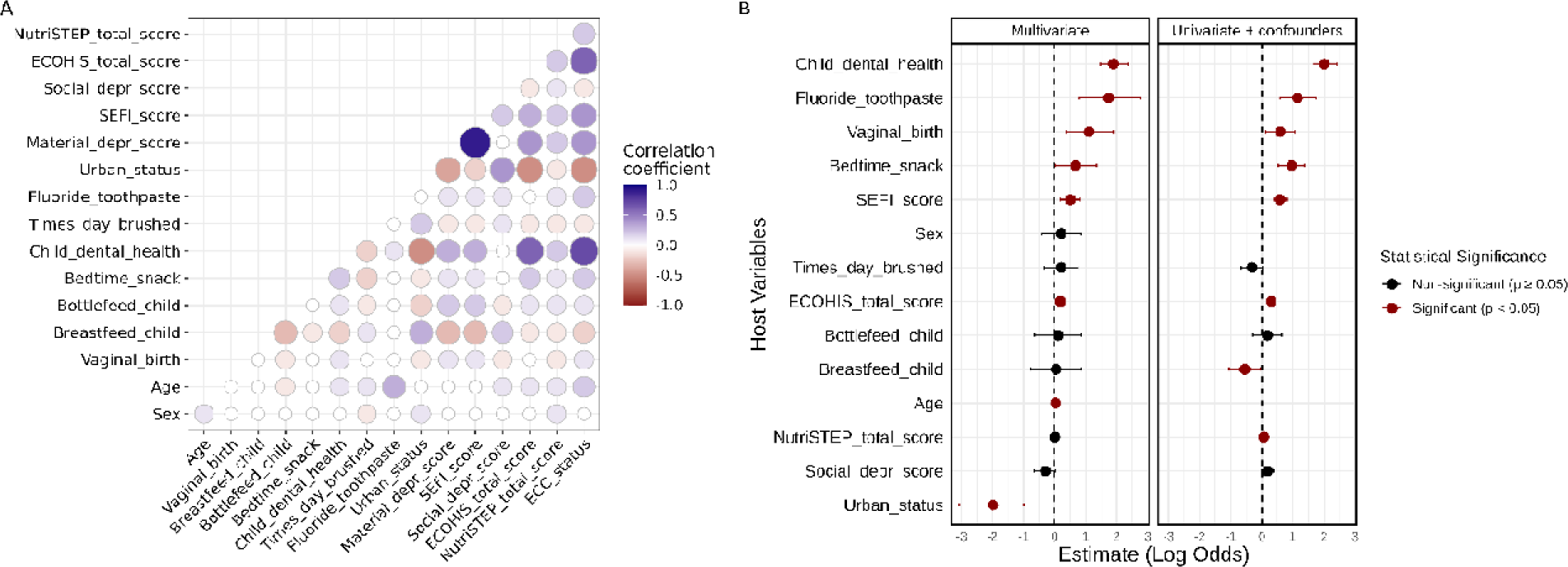
Association between host variables and early childhood caries (A) The correlation heatmap displays the relationships between various socioeconomic and health-related factors. Darker blue circles indicate a stronger positive correlation, while darker red circles represent a stronger negative correlation between the variables. (B) The forest plot illustrates the log odds ratios for the association of each factor with ECC status (right). Factors significantly associated with disease status are highlighted in red, with the horizontal lines representing the 95% confidence intervals. Features with positive log odds values indicate the likelihood of ECC, while negative values signify protective factors against ECC. The left panel shows the estimates for a multivariate model and the right panel shows the result for a univariate model adjusted for age, sex, and residential status.

**Table 2:** Table for participants characteristics.

| **Variable** | **N** | **CF, N = 226<sup>1</sup>** | **ECC, N = 312<sup>1</sup>** | **p-value<sup>2</sup>** |
| --- | --- | --- | --- | --- |
| **Sex** | 538 |  |  | >0.9 |
| Female (0) |  | 114 (50%) | 157 (50%) |  |
| Male (1) |  | 112 (50%) | 155 (50%) |  |
| **Age** | 538 | 43 (28, 55) | 49 (36, 58) | <0.001 |
| **Vaginal_birth** | 530 |  |  | 0.002 |
| No (0) |  | 72 (32%) | 62 (20%) |  |
| Yes (1) |  | 152 (68%) | 244 (80%) |  |
| **Breastfeed_child** | 530 |  |  | <0.001 |
| No (0) |  | 40 (18%) | 113 (37%) |  |
| Yes (1) |  | 184 (82%) | 193 (63%) |  |
| **Bottlefeed_child** | 531 |  |  | 0.012 |
| No (0) |  | 64 (29%) | 59 (19%) |  |
| Yes (1) |  | 160 (71%) | 248 (81%) |  |
| **Bedtime_snack** | 536 |  |  | <0.001 |
| No (0) |  | 143 (64%) | 127 (41%) |  |
| Yes (1) |  | 81 (36%) | 185 (59%) |  |
| **Child_dental_health** | 534 |  |  | <0.001 |
| Very good (1) |  | 118 (53%) | 12 (3.8%) |  |
| Good (2) |  | 91 (41%) | 64 (21%) |  |
| Fair (3) |  | 12 (5.4%) | 113 (36%) |  |
| Poor (4) |  | 1 (0.5%) | 108 (35%) |  |
| Very poor (5) |  | 0 (0%) | 15 (4.8%) |  |
| **Times_day_brushed** | 536 |  |  | 0.003 |
| Never (1) |  | 2 (0.9%) | 2 (0.6%) |  |
| Sometimes (2) |  | 5 (2.2%) | 29 (9.3%) |  |
| Once daily (3) |  | 85 (38%) | 130 (42%) |  |
| Twice daily (4) |  | 129 (58%) | 147 (47%) |  |
| >2 time daily (5) |  | 3 (1.3%) | 4 (1.3%) |  |
| **Fluoride_toothpaste** | 492 |  |  | <0.001 |
| No (0) |  | 63 (30%) | 38 (14%) |  |
| Yes (1) |  | 148 (70%) | 243 (86%) |  |
| **Urban_status** | 534 |  |  | <0.001 |
| Rural (0) |  | 8 (3.6%) | 156 (50%) |  |
| Urban (1) |  | 216 (96%) | 154 (50%) |  |
| **Material_depr_score** | 498 | -0.10 (-0.69, 0.44) | 0.75 (0.06, 1.72) | <0.001 |
| **SEFI_score** | 498 | -0.06 (-0.67, 0.71) | 0.87 (0.04, 1.89) | <0.001 |
| **Social_depr_score** | 498 | 0.17 (-0.36, 1.01) | -0.01 (-0.57, 0.63) | 0.003 |
| **ECOHIS_total_score** | 454 | 0 (0, 1) | 6 (2, 12) | <0.001 |
| **NutriSTEP_total_score** | 455 | 20 (16, 24) | 23 (19, 27) | <0.001 |
| N, Number of non-missing entries for each variable<br>CF, Caries-free<br>ECC, Early childhood caries<br><sup>1</sup>n (%); Median (IQR)<br><sup>2</sup>Pearson's Chi-squared test; Wilcoxon rank sum test; Fisher's exact test |  |  |  |  |

### Microbial diversity and host variables

To assess the impact of each host variable on the dental plaque microbiome, we examined the Spearman correlation between each host variable and Shannon diversity, a measure of alpha diversity that explains diversity within a sample. For bacteria, three variables, SEFI score, frequency of tooth brushing, and breastfeeding status, had a significant impact on Shannon diversity (**Figure 5A**). For fungi, the variables significantly associated with Shannon diversity index were SEFI score, ECC status, and ECOHIS score (**Figure 5A**). The observed relationship between Shannon diversity for bacteria and fungi due to the host variables demonstrated mostly contrasting trends; variables positively correlated with bacterial diversity were inversely correlated with fungal diversity, and the reverse was also true.

**Figure 5:**
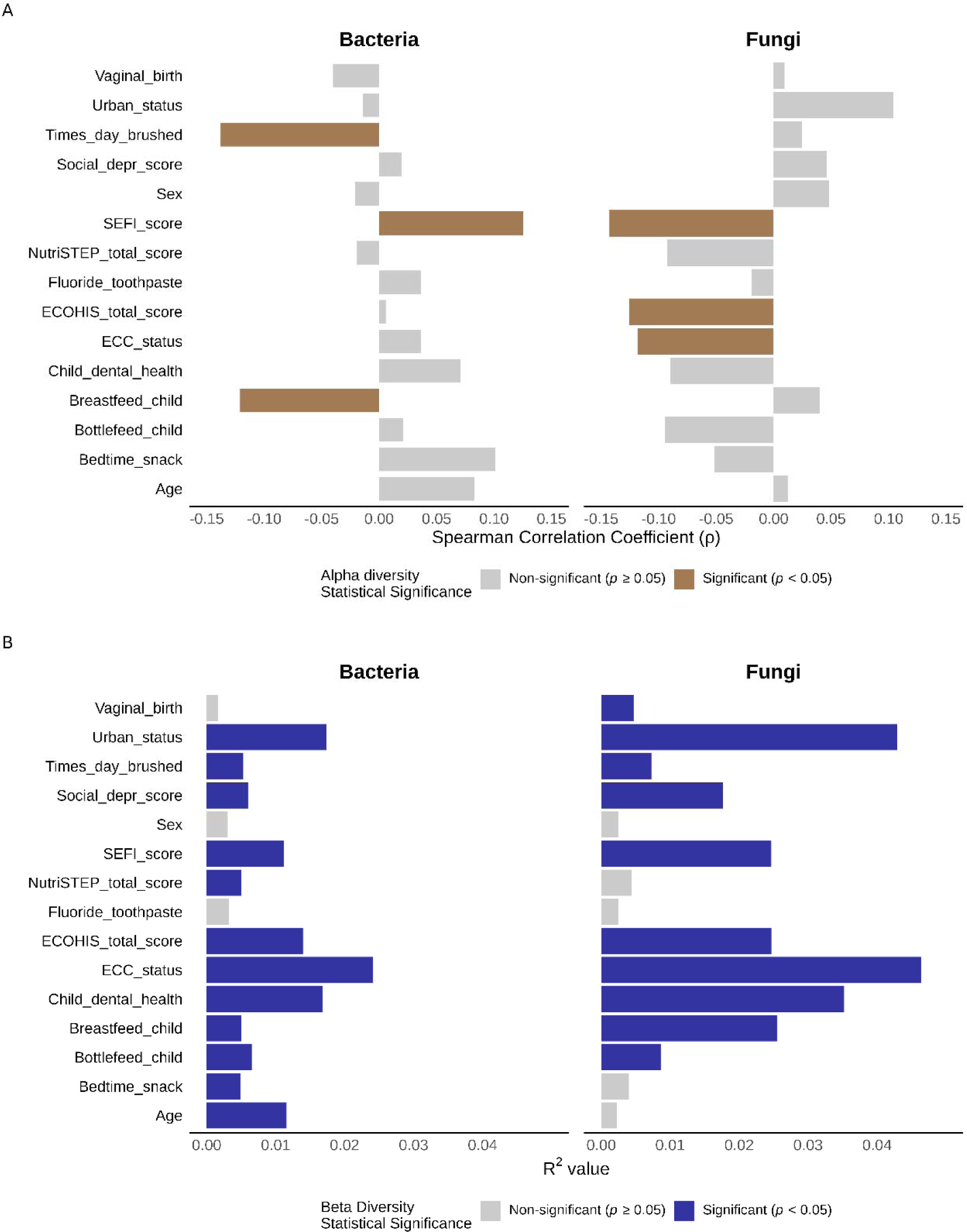
Association of the host variables with diversity metrics (A) Host variables and their Spearman correlation coefficient (ρ) values for alpha diversity, assessed using Shannon’s metric in bacteria and fungi. (B) Beta diversity analyses of host variables via PERMANOVA, with R² values indicating variance explained in bacterial and fungal communities. The colored bar in each plot represents the statistical significance at *p*<0.05.

In the beta diversity analysis, PERMANOVA, where we tested a multivariable model using Bray-Curtis distance as the response variable, identified most of the variables as significant, with the exceptions of sex and use of Fluoride toothpaste for both bacterial and fungal samples, vaginal birth in bacterial samples, and age, NutriSTEP score, and bedtime snacking for fungal samples (**Figure 5B**). However, the proportion of variance (R^2^) explained by the host variables was higher in the fungal microbiome than in the bacterial microbiome.

### Association between microbiome and host variables

The associations of different species-level taxa with host variables are shown in **Figure 6**. The majority of microbial species associated with at least one host variable were linked to ECC status, exhibiting a similar pattern to species influenced by child dental health. After controlling for age, sex, and urban status, the mode of childbirth, NutriSTEP score, and fluoride toothpaste use showed no association with any species. While other variables related to eating habits such as breastfed child, bottle-fed child, and bedtime snacking habit were associated with not more than three species. *S. mutans*, which also showed the most prominent association with ECC status, was also found to be strongly associated with child dental health, SEFI score, and ECOHIS score. *Prevotella* and *Alloprevotella* (formerly classified as Prevotella) species were among the most prevalent in ECC-associated species, for instance, *Prevotella* species *P. salivae, P. oulorum, P. denticola*, *P. nigrescens, P. veroralis*, and *Alloprevotella* sp. HMT-308.

**Figure 6:**
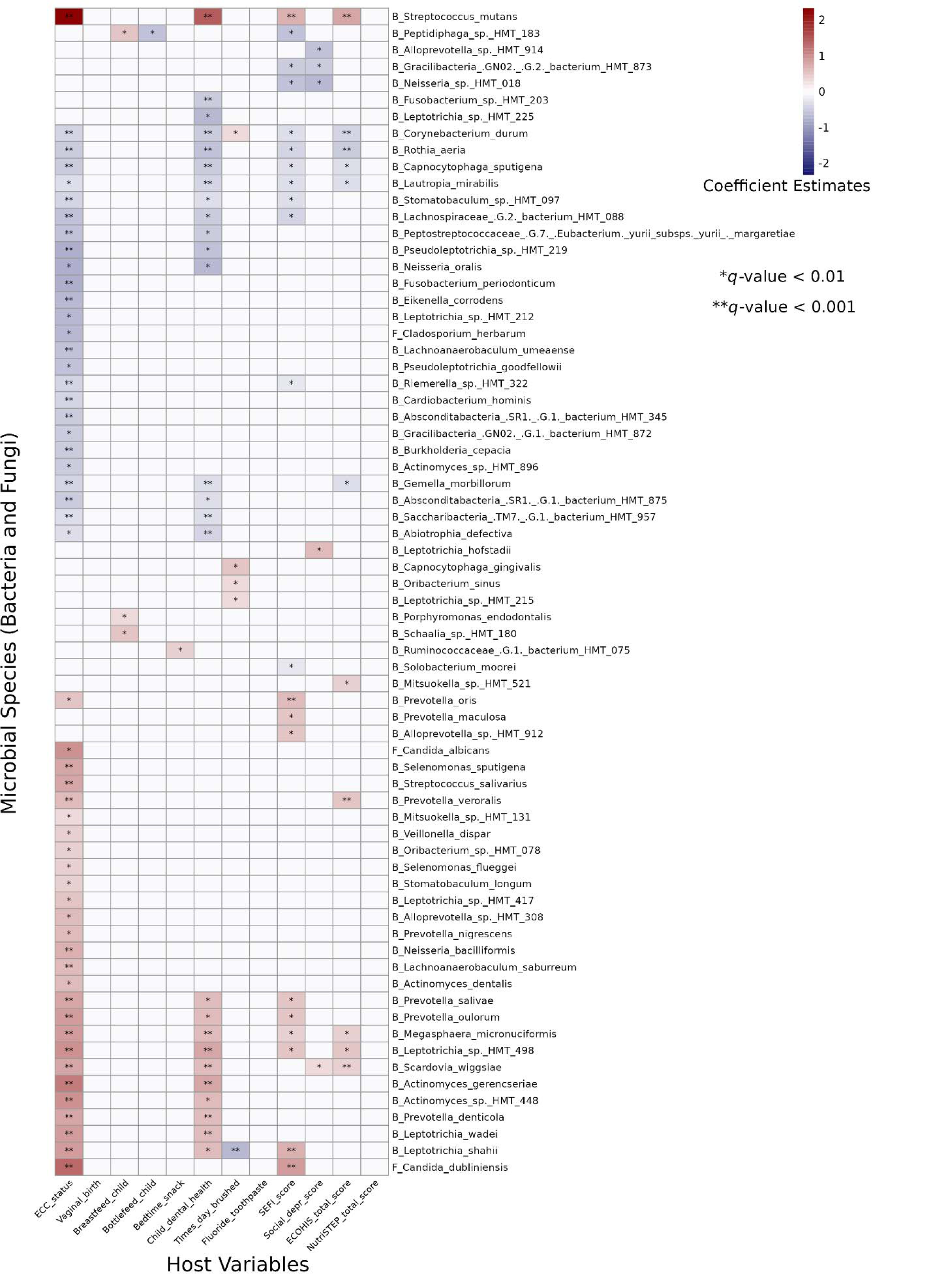
Associations between microbial features and various socioeconomic factors. The coefficient estimate values are after adjusting for age, sex and rural/urban status using MaAsLin2 method for differential abundance analysis. Features are categorized as B_ for bacterial and F_ for fungal. Significance levels are denoted as: * *q*-value < 0.01 and ** *q*-value < 0.001, where *q*-value = Benjamini-Hochberg adjusted *p*-value.

*Alloprevotella* sp. HMT-912, *P. maculosa,* and *P. oris* were unique in that they were positively associated only with socioeconomic variables, SEFI score and social deprivation scores. Many species strongly associated with the SEFI score were also strongly associated with ECC status, even after adjusting for place of residence, suggesting that ECC is highly linked to socioeconomic conditions. This further emphasizes that socioeconomic factors are associated with both caries-causing and caries-protective species.

## DISCUSSION

This study is the first large-scale investigation into how socioeconomic and behavioral factors are associated with the diversity and composition of the dental plaque microbiome in the context of caries and CF conditions in the Canadian preschool population. Previous research on the factors shaping the oral microbiome has primarily focused on ECC status, whereas other host variables included in this study remain largely underexplored. Our research found that socioeconomic status and place of residence (rural or urban), along with certain behavioral factors such as bedtime snacking habits, are associated with ECC, as well as with variations in plaque microbial diversity (**Figure 4B and Figure 5**). Additionally, we identified specific taxonomic differences attributable to these factors (**Figure 6**). Furthermore, we identified potential shifts in the interkingdom interactions between bacteria and fungi in the presence and absence of ECC (**Figure 2**).

The differences in oral microbiome composition were very prominent in fungi, as the two most common species, *C. dubliniensis* and *C. albicans*, occupied close to 75% abundance in ECC, whereas their abundances were very low in children who were CF. In contrast, the taxonomic changes in the bacterial data were more subtle. Fungal data also showed significantly higher Shannon and Simpson diversities in samples from CF children compared to ECC; however, these differences were not significant for bacterial composition. This indicates that while the overall number of species and their distribution within groups may be similar, as suggested by alpha diversity, the specific species present or their relative abundances across samples vary between the groups. For beta diversity, both fungal and bacterial data showed a similar pattern, where differences between the CF and ECC groups were significant but had a low effect on overall differences. The results for alpha diversity are similar to those of our previous publication, where a significant change was observed in Chao1 diversity but not in Shannon diversity in bacteria (de Jesus et al., 2021). A similar trend has been previously observed in the salivary bacteriomes of CF children and those with caries (Grier et al., 2020). Although not significant, the mean Shannon diversity value in our bacteriome was slightly increased in ECC, which is similar to an earlier report on the dental plaque microbiome (Richards et al., 2017). We also found Shannon diversity to be the most commonly used metric for comparing alpha diversity (Alyousef et al., 2023; de Jesus et al., 2021; Grier et al., 2020; Q. Jiang et al., 2018; Richards et al., 2017). Owing to its standardized, relative, and comprehensive analysis, which considers both richness and evenness, we suggest including it in future publications for proper diversity comparison.

Our present study suggests that a species-level analysis is crucial for understanding the etiology of ECC. This importance extends beyond species-specific biochemical transformations, as species belonging to the same genus can exhibit contrasting effects leading to disease pathology. For instance, *N. bacilliformis* was found to be associated with ECC, whereas *Neisseria oralis* was associated with CF status. Similar distinctions were observed with *L. shahii* (ECC) compared to *Leptotrichia* sp. HMT-212 and *Leptotrichia* sp. HMT-225 (CF) and among some *Prevotella* species (**Figure 6**). This conclusion is supported by findings from other studies (S. Jiang et al., 2016; H. Xu et al., 2014). This pattern also applies to the genus *Streptococcus*. While *S. mutans* and *Streptococcus salivarius* were significantly associated with ECC, the genus *Streptococcus* as a whole was not (**Figure S2B**). These results are consistent with those of other studies (S. Jiang et al., 2016; Yang et al., 2023).

Several studies have investigated the interkingdom associations, particularly between bacteria and fungi, in the oral microbiome (Balakrishnan et al., 2021; Sztajer et al., 2014). We found an interaction between *C. dubliniensis* and *N. bacilliformis*, which was exclusively observed in the ECC cohort. However, this novel interaction has not been reported previously. Interestingly, these two species were not found to be positively associated with any other species in the ECC group. Such interkingdom associations within ECC microbiomes derived from dental plaque samples have rarely been reported (de Jesus et al., 2020). Furthermore, most studies on interactions in the oral microbiome are restricted to *C. albicans* and *Streptococci*, specifically, the interactions between *C. albicans* and *S. mutans* and their co-localization in ECC (Du et al., 2022; Montelongo-Jauregui & Lopez-Ribot, 2018). In contrast, our findings suggest that the interactions between *C. albicans* and *S. mutans* are more prominent in the CF group than in the ECC group (**Figure 2**). Our intrakingdom associations also support recent findings where species correlations within the same genus of *Leptotrichia* and *Prevotella* were high (Cho et al., 2023). Furthermore, Cho et al. showed *S. mutans* interactions with *P. salivae* and *Leptotrichia* wadei, which in our results was not direct but appears to be mediated by another species *Mitsuokella* sp. HMT-131. While their *in vivo* study showed that *S. sputigena* interacts with *S. mutans*, a clear correlation was not observed in their metagenomic analysis. Similarly, in our analysis, both *S. sputigena* and *S. mutans* were found to be significantly associated with ECC status, but a correlation between them was not observed. On the other hand, *S. mutans* was found to be significantly correlated with *S. wiggsiae*, supporting a previous study in adolescents’ caries (Eriksson et al., 2018).

ML-based classifiers are instrumental in distinguishing between the two conditions based on microbiome data. Previous research on caries has used ML to identify caries-related biomarkers (Butcher et al., 2022; de Jesus et al., 2021, 2020; Grier et al., 2020). Moreover, these models help to determine the contribution of each species to this classification, which indirectly suggests their empirical importance. We identified RF as the best among the ML algorithms selected in our study. Other studies have also found RF to provide good predictive potential (Butcher et al., 2022; Thomas et al., 2019). In addition to its prediction capacity, RF also provides the importance of the variables used in the model, which aids in identifying the species ranking in outcome prediction. Given the variations in performance shown by other classifiers, hyperparameter spaces, and model interpretability, we recommend using the RF method for such analyses. In our ML analysis for feature importance, in addition to *Streptococcus*, *Candida*, and *Neisseria* species, *S. wiggsiae* emerged as one of the top feature species for classifying ECC and CF samples. The prominence of ECC-associated species in the model can potentially be attributed to the significant shifts observed in these species between CF and ECC conditions. Even though bacterial samples demonstrated superior classification performance for CF and ECC compared to fungal samples, we suggest including both bacterial and fungal species in future studies (**Figure 3A**). This recommendation is based on the observation that fungal species are among the top-ranking features based on their variable importance (**Figure 3B**). The AUROC metric is better suited for perfectly balanced classes. As our samples were not perfectly balanced for disease outcome, we also included the AUPRC metric, which is less affected by class imbalances. The performance remained consistent across both metrics, potentially owing to the large sample size, which is critical for ML, especially when the feature count is nearly 300, as in our case. Our large sample size, supporting the application of ML methods, is evident from the high and similar AUROC and AUPRC values. These limitations and considerations were previously discussed (Grier et al., 2020).

Our study found no significant association between ECC status and sex, consistent with findings from a review of Canadian studies, in which only one of five studies reported sex as a significant variable in ECC (Pierce et al., 2019; Schroth & Cheba, 2007). This finding suggests an equal prevalence of caries among male and female children. We also observed a pattern of differentially abundant species linked to child dental health, similar to that observed in ECC, attributable to a high correlation between these variables. We also identified that a higher SEFI score, which indicates less favorable socioeconomic conditions, is associated with an increased risk of ECC. Additionally, ECC’s association with bedtime snacking habits shows that ECC is influenced by an intricate interplay of biological, behavioral, and socioeconomic factors (Anil & Anand, 2017; Hussein et al., 2017). For differential species analysis, we used age, sex, and place of residence as confounding factors. The influence of age and sex on ECC microbial differences has been shown previously (de Jesus et al., 2020; L. Xu et al., 2018). The variability in differentially abundant species with and without confounding factors is compared in **Figure 6** and **Figure S2A**. The association between ECC and bedtime snacking habits underscores the importance of dental hygiene, especially before bedtime, in preventing caries. Furthermore, the association between fluoride toothpaste and ECC could be attributed to the more prevalent use of fluoride toothpaste by children with existing poor dental health.

A high prevalence of *Prevotella* spp. associated with ECC was observed in our analysis. *Prevotella* species serve as biomarkers for ECC detection and their role as potential periodontal pathogens has been widely recognized (He et al., 2018; Teng et al., 2015; Yang et al., 2023). In contrast, none of the *Lactobacillus* species were identified as ECC-associated. *Lactobacillus* species have been previously found to be associated with dental caries (X. Chen et al., 2020). From our results, *N. oralis* can be considered a potential biomarker for CF status, while the role of *N. bacilliformis* in ECC has been further emphasized by recent studies (Cherkasov et al., 2019; Fakhruddin et al., 2022; E. Lee et al., 2021).

Our findings on fungi are consistent with both our previous study and other research where *C. albicans* and *C. dubliniensis* were the most prevalent species (Al-Ahmad et al., 2016; de Jesus et al., 2020). These two species also appeared as biomarkers in both differential abundance analysis and RF-based classification. The differential abundance results obtained from our analysis were also compared with those derived using an alternative method, LinDA (Zhou et al., 2022). This comparison revealed a high degree of consistency between the outcomes of both the models (data not shown). This consistency strengthens the validity of our results and contributes to a broader understanding of the microbial interactions in ECC.

Our analysis was based on 16S rRNA and ITS sequencing rather than metagenomic data. ITS sequencing is widely accepted for studying fungal composition owing to its reasonable discriminatory power and well-defined reference databases (Conti et al., 2023; Schoch et al., 2012). However, metagenomic sequencing offers additional advantages, as it enhances strain-level resolution, which is somewhat challenging with 16S rRNA and ITS sequencing. We also recognize the value of longitudinal studies and matching participants by sex, age, and socioeconomic status to capture temporal differences, while reducing confounding effects. The lack of species-level information for fungi in some samples, attributed to a low number of reads, may have resulted from insufficient fungal DNA in the original samples, owing to low fungal biomass. Additionally, the UNITE database, commonly used for fungal taxonomic assignments, does not offer as high taxonomic resolution as bacterial databases such as HOMD and Silva (Nilsson et al., 2016; Quast et al., 2013). In the future, enhancing the understanding of ECC could be achieved through longitudinal birth cohort studies. Such an analysis would yield more specific differences and capture dynamic changes over time, while effectively controlling for numerous confounding variables. While many ECC associated species and interactions identified in our analysis corroborated with previous studies, these findings may not be generalized across different populations due to the variations identified by place of residence and socioeconomic conditions. Furthermore, the biases introduced by class imbalance for variables selected in the study, sample processing and sequencing at different time points are also inevitable in microbiome studies, however, some biases were minimized using centered log ratio transformation. Some studies have highlighted the association between host taste genetics and the incidence of caries, including ECC (de Jesus et al., 2022; Orlova et al., 2022).

Therefore, the inclusion of genetic data, especially in the field of taste genetics, may help to identify the role of nutrition and food intake in the susceptibility to ECC. Furthermore, multimodal machine learning tools can identify ECC contributors to the microbiome, host behaviors, socioeconomic status, and genetic components.

In summary, our study investigated the impact of dental plaque microbiome, socioeconomic, and behavioral factors on the presence of ECC. We reported several novel interactions between the bacteriome and mycobiome in the ECC and CF groups. This includes interkingdom interactions between *C. dubliniensis* and *N. bacilliformis*, and *C. albicans* and *C. durum*. Our study identified key species associated with ECC, including *S. mutans*, *C. dubliniensis*, *N. bacilliformis*, among others. This analysis was facilitated using microbiome association analysis methods and ML models, made possible by inclusion of mycobiome sequencing and a large sample size. This study provides empirical evidence linking socioeconomic and behavioral factors, such as SEFI score and bedtime snacking habits, to ECC and offers new insights into the contributors to ECC. Our research will aid oral healthcare providers in the development and implementation of targeted intervention strategies for ECC based on the specific host variables included in the study.

## DATA AND CODE AVAILABILITY

The data used in this study and additional information required to reanalyze the data reported in this paper is available from the lead contact upon request.

## ACKNOWLEDGMENTS

This study was funded by the Canadian Institutes of Health Research (CIHR) through an operating grant PJT-159731. Both MWK and VCJ received GETS funding from the University of Manitoba. Additionally, MWK was supported by the Natural Sciences and Engineering Research Council of Canada (NSERC) through the VADA grant. We thank the participants, the parents and caregivers, and the Misericordia Health Centre. The authors would also like to express their gratitude to Drs. Nisha Singh, Ryan Cunnington, and Ankita Vaishampayan of the Chelikani Lab for their invaluable assistance and logistical support.

## AUTHOR CONTRIBUTIONS

- MWK, RJS, PH, and PC conceived the study.
- MWK, VCJ, PH, and PC contributed to study design, data analysis, interpretation, and writing of the manuscript.
- MWK, VCJ, BAM, VL, and RJS contributed to data acquisition.
- BAM, VL, and SS contributed to data preparation and analysis.
- RJS contributed to the study design, data interpretation, and manuscript writing.
- PH, RJS, and PC contributed to the funding acquisition. All authors have contributed to the manuscript and approved the submitted version.

## STAR METHODS

### KEY RESOURCES TABLE

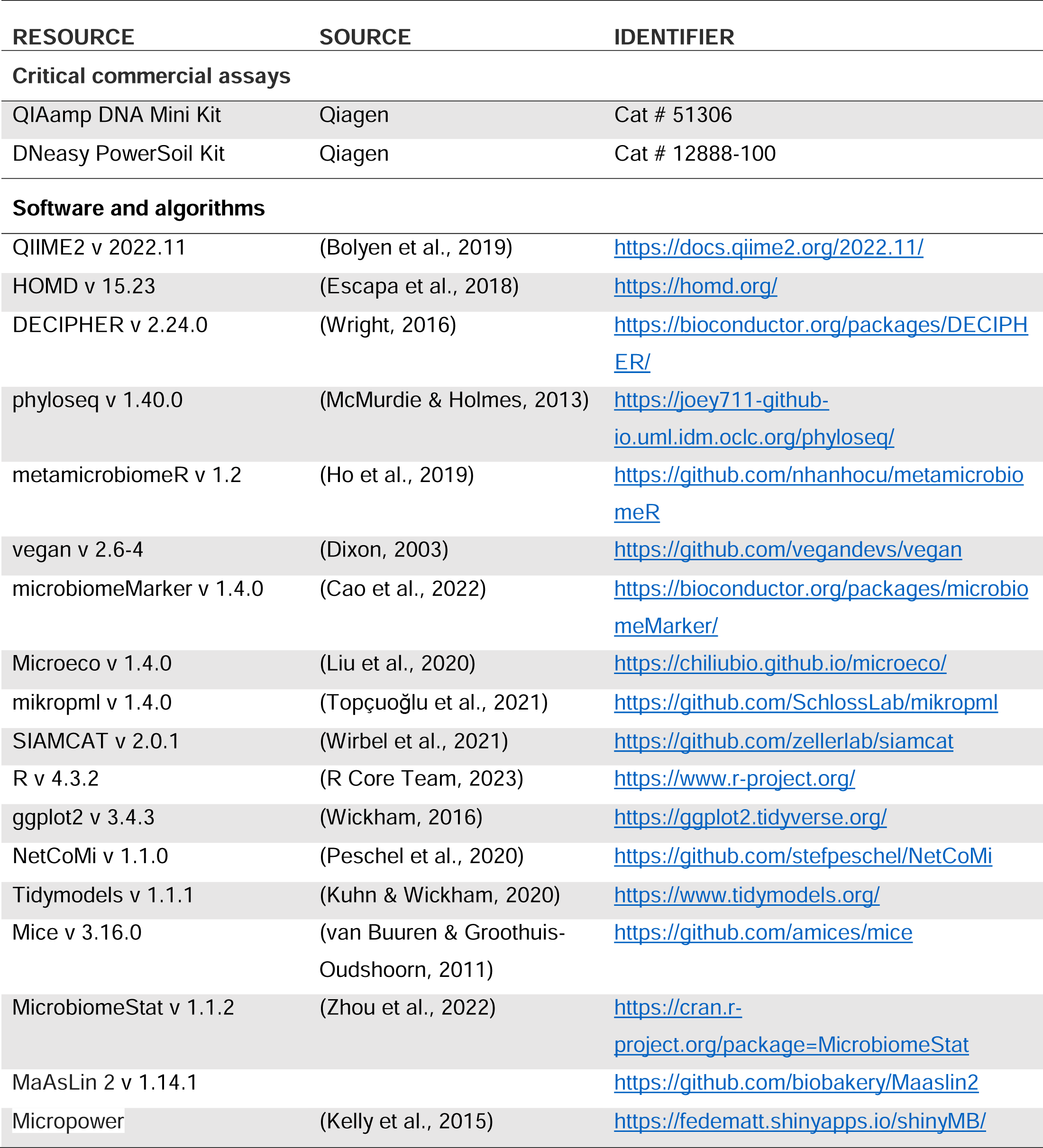

---

**Figure S1:**
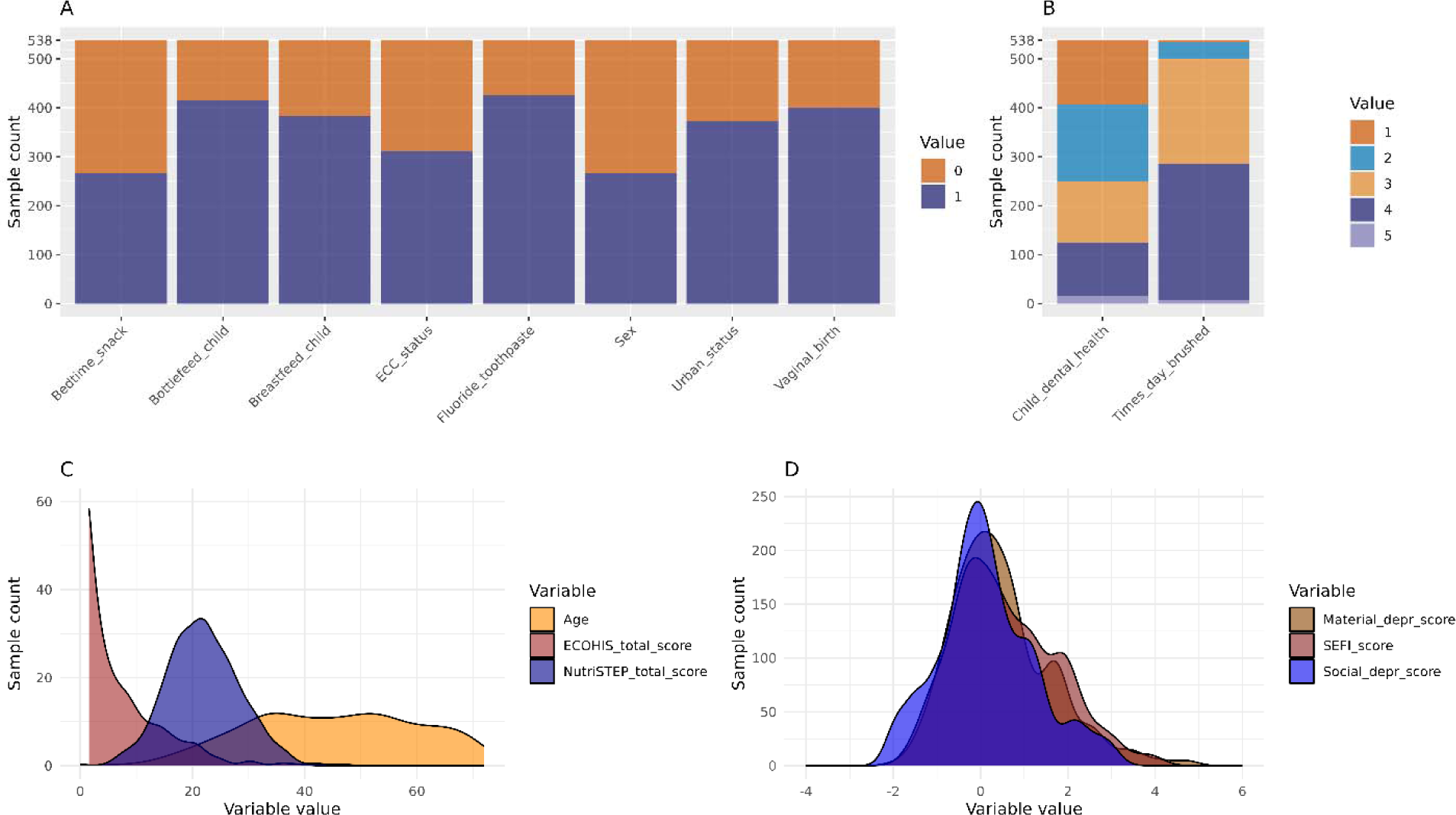
Exploratory data analysis for the distribution of host variables used as socioeconomic and behavioral factors after imputation (A) Binary variables. (B) Ordinal Variables. (C) Continuous variables for age, ECOHIS AND NutriSTEP score. (D) Continuous variables for socioeconomic factors. The detail of these variables is provided in Table 1 in the main article.

**Figure S2:**
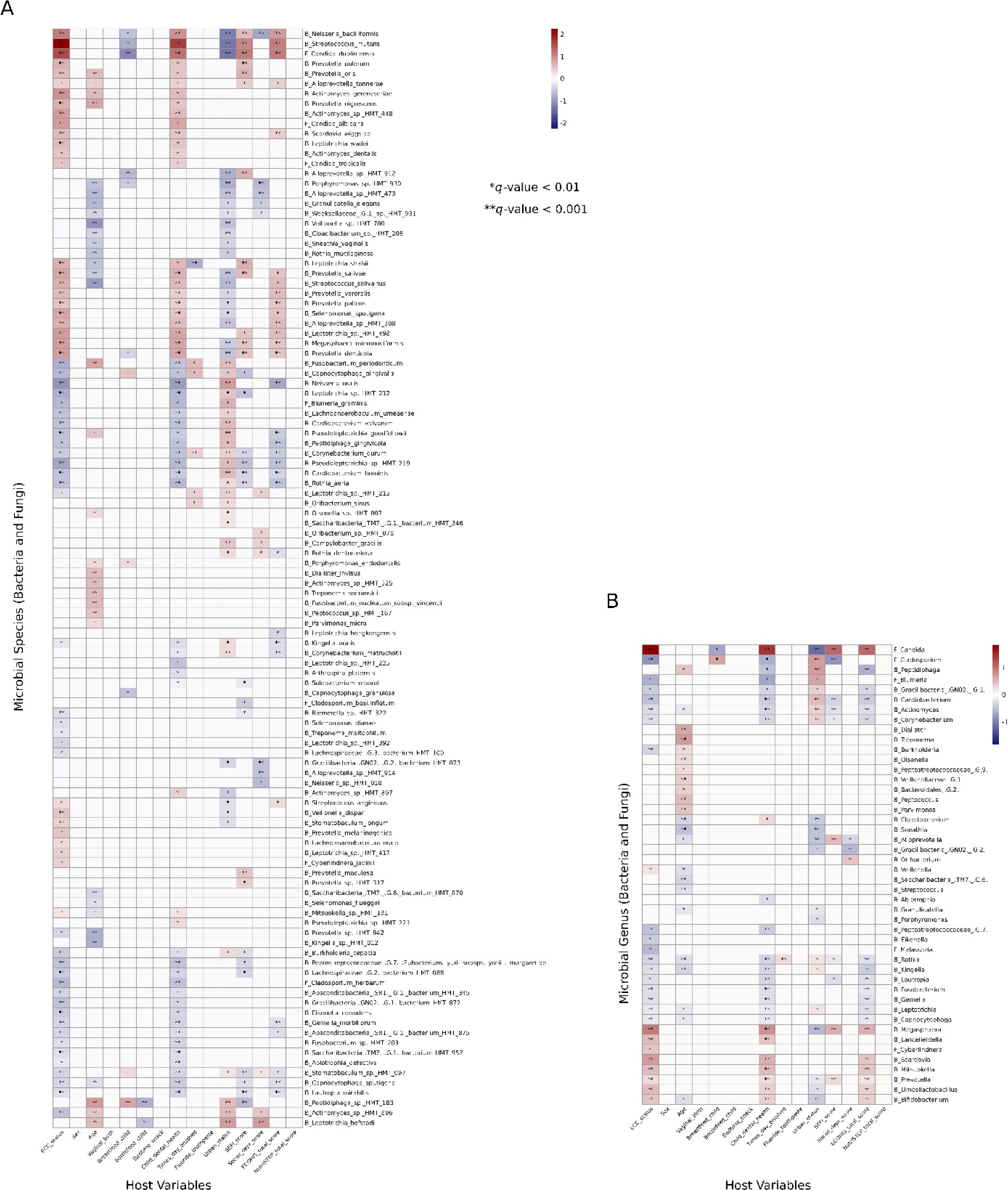
Association between microbial features and various socioeconomic factors (A) Species level. (B) Genus level. The coefficient estimates values from MaAsLin2 differential abundance analysis method without adjusting for any confounder. Features are categorized as B_ for bacterial and F_ for fungal. Significance levels are denoted as: * q-value < 0.01 and ** q- value < 0.001, where q-value = Benjamini-Hochberg adjusted p-value.

**Table S1:** STROBE Statement—Checklist of items that should be included in reports of *cross-sectional studies*.

|  | Item No | Recommendation | Page No |
| --- | --- | --- | --- |
| Title and abstract | 1 | (a) Indicate the study's design with a commonly used term in the title or the abstract | 1 |
|  |  | (b) Provide in the abstract an informative and balanced summary of what was done and what was found | 2 |
| Introduction |  |  |  |
| Background/rationale | 2 | Explain the scientific background and rationale for the investigation being reported | 3 |
| Objectives | 3 | State specific objectives, including any prespecified hypotheses | 3-4 |
| Methods |  |  |  |
| Study design | 4 | Present key elements of study design early in the paper | 4 |
| Setting | 5 | Describe the setting, locations, and relevant dates, including periods of recruitment, exposure, follow-up, and data collection | 4 |
| Participants | 6 | (a) Give the eligibility criteria, and the sources and methods of selection of participants | 4 |
| Variables | 7 | Clearly define all outcomes, exposures, predictors, potential confounders, and effect modifiers. Give diagnostic criteria, if applicable | 4-6 |
| Data sources/<br>measurement | 8 | For each variable of interest, give sources of data and details of methods of assessment (measurement). Describe comparability of assessment methods if there is more than one group | 4-6 |
| Bias | 9 | Describe any efforts to address potential sources of bias | 12 |
| Study size | 10 | Explain how the study size was arrived at | 4 |
| Quantitative variables | 11 | Explain how quantitative variables were handled in the analyses. If applicable, describe which groupings were chosen and why | 5-6 |
| Statistical methods | 12 | (a) Describe all statistical methods, including those used to control for confounding | 5-6 |
|  |  | (b) Describe any methods used to examine subgroups and interactions | NA |
|  |  | (c) Explain how missing data were addressed | 6 |
|  |  | (d) If applicable, describe analytical methods taking account of sampling strategy | 5-6 |
|  |  | (e) Describe any sensitivity analyses | NA |
| Results |  |  |  |
| Participants | 13 | (a) Report numbers of individuals at each stage of study—eg numbers potentially eligible, examined for eligibility, confirmed eligible, included in the study, completing follow-up, and analysed | 4 |
|  |  | (b) Give reasons for non-participation at each stage | 4 |
|  |  | (c) Consider use of a flow diagram | NA |
| Descriptive data | 14 | (a) Give characteristics of study participants (eg demographic, clinical, social) and information on exposures and potential confounders | 8 |
|  |  | (b) Indicate number of participants with missing data for each variable of interest | 16-17 |
| Outcome data | 15 | Report numbers of outcome events or summary measures | 8, 16-17 |
| Main results | 16 | (a) Give unadjusted estimates and, if applicable, confounder-adjusted estimates and their precision (eg, 95% confidence interval). Make clear which confounders were adjusted for and why they were included | 8 |
|  |  | (b) Report category boundaries when continuous variables were categorized | NA |
|  |  | (c) If relevant, consider translating estimates of relative risk into absolute risk for a meaningful time period | NA |
| Other analyses | 17 | Report other analyses done—eg analyses of subgroups and interactions, and sensitivity analyses | NA |
| **Discussion** |  |  |  |
| Key results | 18 | Summarise key results with reference to study objectives | 9,12 |
| Limitations | 19 | Discuss limitations of the study, taking into account sources of potential bias or imprecision. Discuss both direction and magnitude of any potential bias | 12 |
| Interpretation | 20 | Give a cautious overall interpretation of results considering objectives, limitations, multiplicity of analyses, results from similar studies, and other relevant evidence | 9-12 |
| Generalisability | 21 | Discuss the generalisability (external validity) of the study results | 12 |
| **Other information** |  |  |  |
| Funding | 22 | Give the source of funding and the role of the funders for the present study and, if applicable, for the original study on which the present article is based | 13 |

